# Accurate Protein-Protein Interactions Modeling through Physics-informed Geometric Invariant Learning

**DOI:** 10.1101/2025.07.01.662544

**Authors:** Jiahua Rao, Deqin Liu, Xiaolong Zhou, Qianmu Yuan, Wentao Wei, Wei Lu, Jixian Zhang, Yu Rong, Yuedong Yang, Shuangjia Zheng

## Abstract

AlphaFold has set a new standard for predicting protein structures from primary sequences; however, it faces challenges with protein complexes across species, engineered proteins, and antigen-antibody, where co-evolutionary signals may be sparse or missing. Herein, we present ProTact, a SE(3)-invariant geometric graph neural network that integrates physics-informed geometric complementarity and trigonometric constraints as robust inductive biases to enhance protein-protein contact predictions. ProTact is applicable to both experimental and predicted monomer structures and utilizes a modulated key point matching algorithm to approximate accurate docking poses. Experimental evaluations demonstrate that ProTact consistently outperforms state-of-the-art sequence-based and structure-based methods on benchmark datasets, achieving notable relative improvements of 31.63% in Precision@10 for CASP 13 and 14 targets, and 31.94% for DIPS-Plus datasets. Moreover, when combined with AlphaFold3 as re-scoring functions, ProTact surpasses its default confidence scores, offering over 30.48% improvements in the low-MSA contexts. We anticipate that the proposed framework will advance our understanding of protein interactions, functions, and design.

## 1 Introduction

Protein-protein interactions (PPIs) are fundamental to numerous biological processes, such as signal transduction, enzymatic reactions, and cellular transport [1–3]. Understanding the structure of these complexes is crucial for developing new therapeutic strategies and elucidating the molecular basis of diseases [4]. Although powerful experimental methods, including X-ray crystallography, nuclear magnetic resonance (NMR) spectroscopy, and cryo-electron microscopy (cryo-EM), are widely used, they can be expensive and technically demanding [5–7]. Consequently, there is a pressing need for efficient computational methods to model the three-dimensional structures of protein complexes [8–11].

Recent advances in deep learning, such as AlphaFold [8, 11], RosettaFold [9, 12], and HelixFold3 [13], have demonstrated unprecedented performance in predicting protein-protein complex structures, revolutionizing the field of structural biology. Despite their impressive power, researchers have emphasized the importance of protein homologs and multiple sequence alignments (MSAs) within frameworks. Co-evolutionary contacts are the foundation for most protein structure prediction methods. Although MSAs provide critical evolutionary information, they can result in less reliable predictions for proteins with insufficient evolutionary data, such as those from viruses, understudied organisms, antibodies [14], synthetic proteins, or those containing clinically relevant mutations [15].

One solution to this problem is to incorporate contact restraints into protein-protein docking. These contact restraints provide crucial co-evolutionary information about the complex interface and can often be acquired relatively easily and at low cost [16]. Recent advances in docking algorithms that consider restraints have shown promising results in improving the accuracy of protein complex predictions, such as ZDOCK [17], HDOCK [18], AlphaLink [19], and ColabDock [16]. Furthermore, the accuracy of contact restraints can be utilized as a custom scoring function to refine the final predictions of methods like AlphaFold [11] and RosettaFold [9]. Developing robust computational methods that can accurately predict inter-chain contacts is therefore crucial.

Despite significant progress in predicting intra-chain contacts or distances [20–25], the performance of these methods still relies heavily on homologies available from MSAs and paired MSAs (interlogs) [26–28]. These methods struggle to effectively integrate many-body interactions and molecular surface features and to adequately model geometric complementarity and invariance. Addressing these challenges is crucial for advancing the predictive capabilities for protein-protein inter-chain contacts, especially when leveraging scarce MSA information.

In this work, we present ProTact, a SE(3)-invariant geometric graph neural network that integrates physics-informed geometric complementarity and trigonometric constraints as robust inductive biases to strengthen co-evolutionary information and improve the accuracy of inter-protein contact predictions. ProTact is designed to capture many-body interactions in inter-chain with surface vector features, which are essential for leveraging the limited information provided by MSAs in co-evolutionary analysis. To incorporate geometric complementarity and invariance, a novel SE(3)-invariant geometric-aware architecture is introduced to process surface messages between two rigid entities.

ProTact consistently outperforms state-of-the-art sequence-based and structure-based methods on benchmark datasets, achieving a 31.63% relative improvement in Precision@10 for CASP13&14 targets and a 31.94% relative gain for DIPS-Plus sets. These results demonstrate the effectiveness of ProTact in enhancing the accuracy of inter-protein contact predictions. We have also shown that the ProTact framework is not limited to experimental structures. Our experiments demonstrate how ProTact can effectively predict contacts for proteins without available experimental structures, working in conjunction with structure prediction methods. Furthermore, ProTact is applicable to employ a modulated key point matching algorithm to approximate accurate docking poses, and its superiority in docking accuracy is validated using the Kabsch algorithm.

To further showcase the potency of ProTact, we demonstrate its ability to enhance the performance of existing docking methods. Experiments show that re-ranking AlphaFold3 predictions using ProTact results in structures that out-perform those obtained with AlphaFold3 alone. Additionally, through transfer learning, we demonstrate that ProTact has potential applications in predicting antigen-antibody complexes, especially in challenging cases with scarce MSA information. These results prove that by integrating robust inductive biases and advanced geometric and surface features, ProTact paves the way for more accurate and scalable prediction methods, potentially transforming our understanding and manipulation of protein complexes in both basic research and therapeutic applications.

## 2 Results

### 2.1 Overview of the ProTact Framework

Figure 1 illustrates our proposed model, ProTact, aimed at predicting inter-chain contacts in protein-protein interactions (PPIs). The architecture comprises two main components: a SE(3)-invariant encoder (SEInvi) and a trigonometry module (TrigModule). The SE(3)-invariant encoder (SEInvi) transforms protein structure and sequence information into rotation- and translation-invariant representations. It uses spherical harmonics encoders to convert pairwise representations into node features, followed by a convolution layer that updates these features based on the edge features of neighboring nodes. This ensures that the resulting node features are independent of the coordinate system, making the model robust to different orientations and positions of the protein. The trigonometry module (TrigModule) integrates trigonometric constraints and molecular surface features to capture critical points in the embedding space and model complex many-body effects. Inspired by works such as AlphaFold [8, 11] and our previous work TANKBind [29], the trigonometry network uses geometric graph neural networks (GNNs) to learn structural representations of the input protein, reflecting the spatial organization and topological neighborhood of the protein structures. The GNNs in TrigModule update the node features by considering both the edge features (which encode residue contacts) and the surface vector features. This integration allows the model to capture the intricate spatial relationships and physical properties of the residues, thereby enhancing the prediction of inter-chain contacts. These learned representations are then combined to predict the inter-chain contacts. The residue surface features and the modeling of many-body effects enable ProTact to accurately predict protein-protein docking interfaces, even with limited Multiple Sequence Alignment (MSA) information. The ProTact framework is not limited to experimental structures. Thanks to the TrigModule’s ability to effectively capture interaction information, ProTact can accurately predict inter-chain contacts for proteins without available experimental structures, working in conjunction with structure prediction methods. Moreover, ProTact can enhance the performance of other docking methods, such as HDock [18] and AlphaFold3 [11], by incorporating its predicted contacts or re-ranking their predictions.

**Fig. 1.**
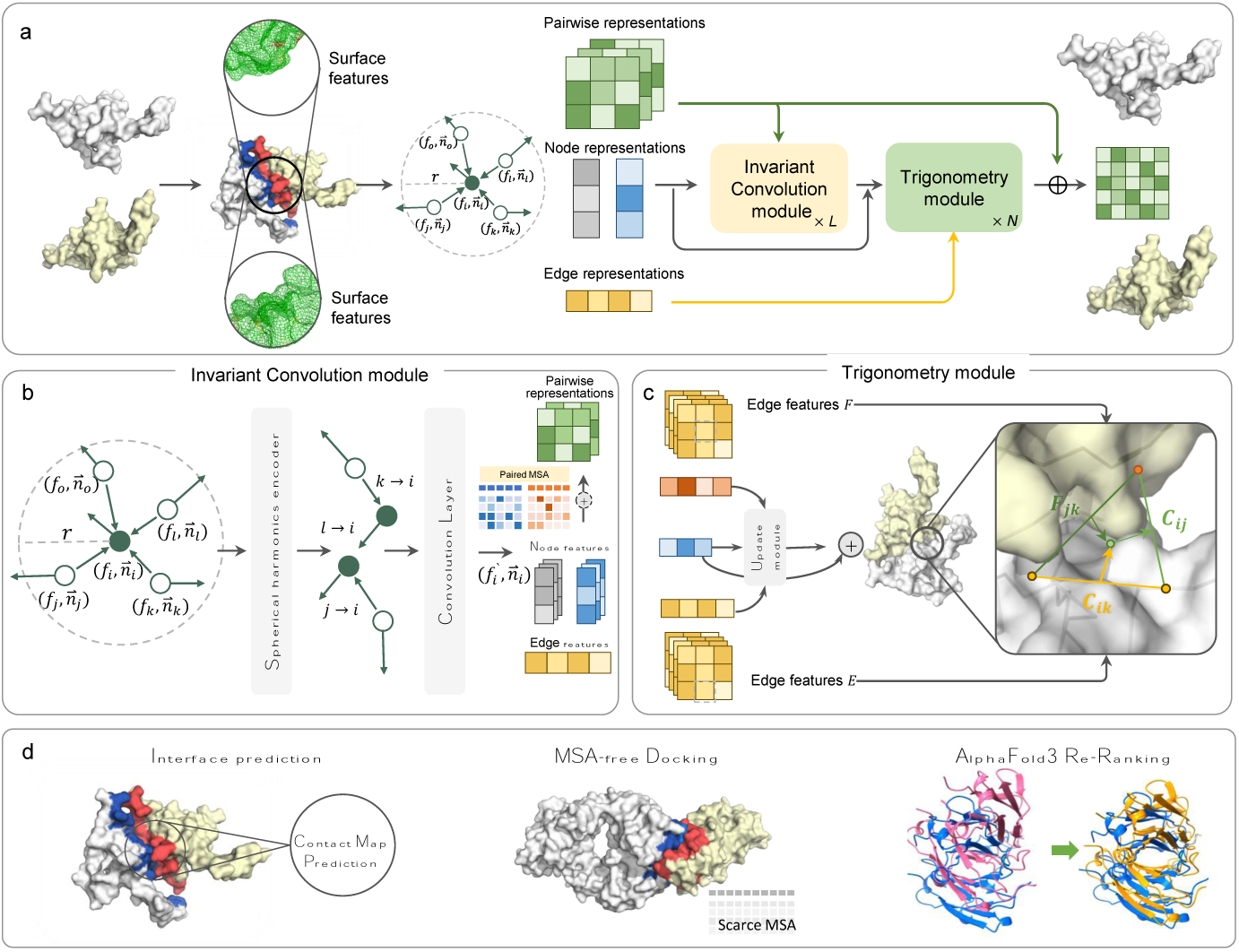
(a) Overview of the contacts prediction and docking workflow. (b) SE(3)-invariant encoder (SEInvi) architecture. The SEInvi module update the input features from equivariance to invariance by convoluting spherical harmonics of the dihedral angle. (c) The trigonometry module (TrigModule) ensures that the interaction between two residues from different proteins depends on all edges in both proteins, considering trigonometry and physical constraints. (d) ProTact demonstrates its versatility by being applicable to a range of prediction tasks, including interface prediction, and docking with and without MSA information, and re-ranking docking predictions.

### 2.2 ProTact achieves state-of-the-art performance in inter-chain contact prediction

To demonstrate the superiority of ProTact, we followed the settings established by [24] and compared our model with several state-of-the-art predictors, including BIPSPI [21], DeepInteract [24], GLINTER [25], CDPred [22], and DeepInter [30], on the DIPS-Plus [31] and CASP13&14 [32, 33] benchmark datasets. The evaluation metrics used for comparison are average top-k precision (P@k), recall (R@k), and Area Under the Curve (AUC).

As shown in Table 1, our model consistently outperformed all other predictors in terms of precision and recall across various top-k thresholds and AUC on both the DIPS-Plus and CASP13&CASP14 test targets. Specifically, on the DIPS-Plus dataset, our model achieved the highest precision of 0.413, 0.419, and 0.376 for P@10, P@L/10, and P@L/5, respectively. Our model showed an improvement of 31.95%, 41.08%, and 31.01% over the best baseline DeepInter (0.313, 0.297, 0.287) under the same condition. When comparing with AlphaFold-Multimer, ProTact also exhibits clear advantages. For instance, on the DIPS-Plus dataset, ProTact’s P@10 score is 0.58, markedly higher than AlphaFold-Multimer’s 0.357. This result highlights its leading position in the field of protein-protein interaction prediction. For the CASP13&14 heteromeric complexes with largely asymmetric inter-chain geometries, our model reached precision values of 0.437, 0.406, and 0.399 for P@10, P@L/10, and P@L/5, respectively. The interfaces predicted by ProTact also had higher performances than all other methods. These consistent results support our assumption that the integration of many-body geometric complementarity and molecular surface features is beneficial to interfacial contact prediction. Table S1 details the results on both DIPS-Plus and CASP13&14 test targets for homodimers and heterodimers, respectively.

**Table 1.** The average top-k precision and recall metrics for different protein-protein interaction prediction methods on the DIPS-Plus and CASP13&14 test targets. Five predictors are compared: BIPSPI, DeepInteract, GLINTER, CDPred, DeepInter, and ProTact. For each predictor, seven metrics are evaluated: Precision at 10 (P@10), Precision at L/10 (P@L/10), Precision at L/5 (P@L/5), Recall at L (R@L), Recall at L/2 (R@L/2), Recall at L/5 (R@L/5), and Area Under the Curve (AUC). The best and second best results are marked **bold** and underline, respectively.

| DIPS-Plus |  |  |  |  |  |  |  |
| --- | --- | --- | --- | --- | --- | --- | --- |
| Predictor | P@10 | P@L/10 | P@L/5 | R@L | R@L/2 | R@L/5 | AUC |
| BIPSPI | 0.010 | 0.010 | 0.010 | 0.010 | 0.004 | 0.003 | - |
| DeepInteract | 0.206 | 0.207 | 0.182 | 0.158 | 0.098 | 0.046 | 0.90 |
| GLINTER | 0.234 | 0.222 | 0.189 | 0.154 | 0.102 | 0.056 | 0.87 |
| CDPred<sup>a</sup> | 0.154 | 0.152 | 0.125 | 0.086 | 0.056 | 0.034 | 0.86 |
| DeepInter | 0.313 | 0.297 | 0.287 | <u>0.201</u> | 0.124 | 0.058 | <u>0.92</u> |
| AF-Multimer<sup>b</sup> | <u>0.391</u> | <u>0.387</u> | **0.386** | 0.191 | <u>0.173</u> | <u>0.094</u> | - |
| ProTact | **0.413** | **0.419** | <u>0.376</u> | **0.324** | **0.204** | **0.103** | **0.96** |
| CASP13&14 |  |  |  |  |  |  |  |
| Predictor | P@10 | P@L/10 | P@L/5 | R@L | R@L/2 | R@L/5 | AUC |
| BIPSPI | 0.010 | 0.000 | 0.010 | 0.020 | 0.010 | 0.001 | - |
| DeepInteract | 0.181 | 0.192 | 0.140 | 0.113 | 0.074 | 0.037 | 0.86 |
| GLINTER | 0.213 | 0.203 | 0.167 | 0.102 | 0.066 | 0.038 | 0.82 |
| CDPred<sup>c</sup> | 0.142 | 0.147 | 0.122 | 0.101 | 0.074 | 0.027 | 0.90 |
| DeepInter | <u>0.332</u> | <u>0.338</u> | <u>0.306</u> | <u>0.195</u> | <u>0.127</u> | <u>0.062</u> | <u>0.94</u> |
| ProTact | **0.437** | **0.406** | **0.399** | **0.338** | **0.224** | **0.112** | **0.98** |

<sup>a</sup>CDPred succeed predict only 12 homo and 12 hetero protein complexes.
<sup>b</sup>We selected and ranked the 50 inter-protein residue pairs with the shortest heavy atom distances in each generated protein complex structures as the predicted inter-protein contacts.
<sup>c</sup>CDPred succeed predict only 9 homo and 2 hetero protein complexes.

To ensure a more fair and comprehensive comparison, we further collected the heterodimers and homodimers published between 2023-01-01 and 2023-06-31 in the PDB to create a larger blind test set, including 191 heterodimers and 41 homodimers. Table 2 summarizes the experimental results on this blind test dataset for various predictors, including DeepInteract [24], DeepInter [30], PLMGraph-Inter [34], and ProTact. ProTact achieved the highest Precision at 10, Precision at L/10, and Precision at L/5 for both homodimers and heterodimers, with values of 0.210, 0.213, and 0.190 for homodimers and 0.191, 0.190, and 0.179 for heterodimers, respectively. ProTact also had the highest Recall at L and Recall at L/2 scores for both homodimers (0.071 and 0.045) and heterodimers (0.054 and 0.033). Furthermore, ProTact achieved the highest AUC score of 0.707 for heterodimers and 0.701 for homodimers. Compared to the baseline methods, ProTact demonstrated significant improvements, with average improvements of 39.08%, 39.26%, and 34.92% over PLMGraph-Inter and 93.14%, 92.86%, and 78.00% over DeepInter for Precision at 10, Precision at L/10, and Precision at L/5, respectively. These results underscore the superior performance of ProTact in accurately predicting inter-chain contacts in both homodimers and heterodimers, highlighting its potential as a powerful tool for predicting protein-protein interactions.

**Table 2.** The experiments results of Precision at 10 (P@10), Precision at L/10 (P@L/10), Precision at L/5 (P@L/5), Recall at L (R@L), Recall at L/2 (R@L/2), Recall at L/5 (R@L/5), and Area Under the Curve (AUC) on the blind test dataset for DeepInteract, DeepInter, PLMGraph-Inter, and ProTact.

| Homodimers (41) |  |  |  |  |  |  |
| --- | --- | --- | --- | --- | --- | --- |
| Predictor | P@10 | P@L/10 | P@L/5 | R@L | R@L/2 | AUC |
| DeepInteract | 0.054 | 0.058 | 0.118 | 0.016 | 0.013 | 0.338 |
| DeepInter | 0.046 | 0.042 | 0.051 | 0.024 | 0.013 | 0.656 |
| PLMGraph-Inter | 0.151 | 0.135 | 0.126 | 0.051 | 0.029 | 0.684 |
| ProTact | **0.210** | **0.213** | **0.190** | **0.071** | **0.045** | **0.701** |
| Heterodimers (191) |  |  |  |  |  |  |
| Predictor | P@10 | P@L/10 | P@L/5 | R@L | R@L/2 | AUC |
| DeepInteract | 0.039 | 0.034 | 0.028 | 0.022 | 0.011 | 0.316 |
| DeepInter | 0.102 | 0.104 | 0.100 | 0.034 | 0.018 | 0.621 |
| PLMGraph-Inter | 0.162 | 0.172 | 0.161 | 0.050 | 0.031 | 0.693 |
| ProTact | **0.191** | **0.190** | **0.179** | **0.054** | **0.033** | **0.707** |

One advantage of the ProTact framework is its robustness to structural noise. The rapid advancement of protein structure prediction methods has led to an exponential increase in the availability of structural data for analysis. This scenario highlights the growing need for a generalized computational model to analyze predicted structures. We aimed to demonstrate that ProTact is capable of maintaining superior predictive performance whether using experimental structures or predicted structures. As shown in Table S3, despite a slight decrease in performance when using AlphaFold2-predicted structures, ProTact maintains high accuracy in contact predictions across both the DIPS-Plus and CASP13&14 datasets. Specifically, on the DIPS-Plus Homo dataset, ProTact achieves P@10, P@L/10, and P@L/5 scores of 0.58, 0.57, and 0.52, respectively, when using experimental structures. Even when the input is changed to AlphaFold2-predicted structures, these values remain at a high level with P@10, P@L/10, and P@L/5 scores of 0.45, 0.41, and 0.40, indicating ProTact’s strong tolerance for structural prediction errors.

### 2.3 ProTact is robust and generalizable for inter-chain contact prediction

To investigate the contributions of different components, we conducted an ablation study to validate the effectiveness of ProTact. As shown in Figure 2a, incorporating geometric surface features such as vector-based (vSurf) and angle-transformed (aSurf) variants into the TrigModule improved precision and recall rates. Specifically, the E3NN+vSurf+TrigModule variant achieved higher precision and recall compared to the baseline, supporting our hypothesis that geometric surface features enhance the model’s ability to learn complementarity. Comparing the SEInvi module to the E3NN proposed by DeepInteract [24], we observed significant improvements in prediction performance. Combining SEInvi with ResNet [35] or TrigModule further enhanced performance, with SEInvi + TrigModule achieving the highest scores. The ablation study demonstrated the importance of integrating SEInvi and TrigModule, allowing for more accurate and effective prediction of protein contact maps, surpassing previous models and individual components.

**Fig. 2.**
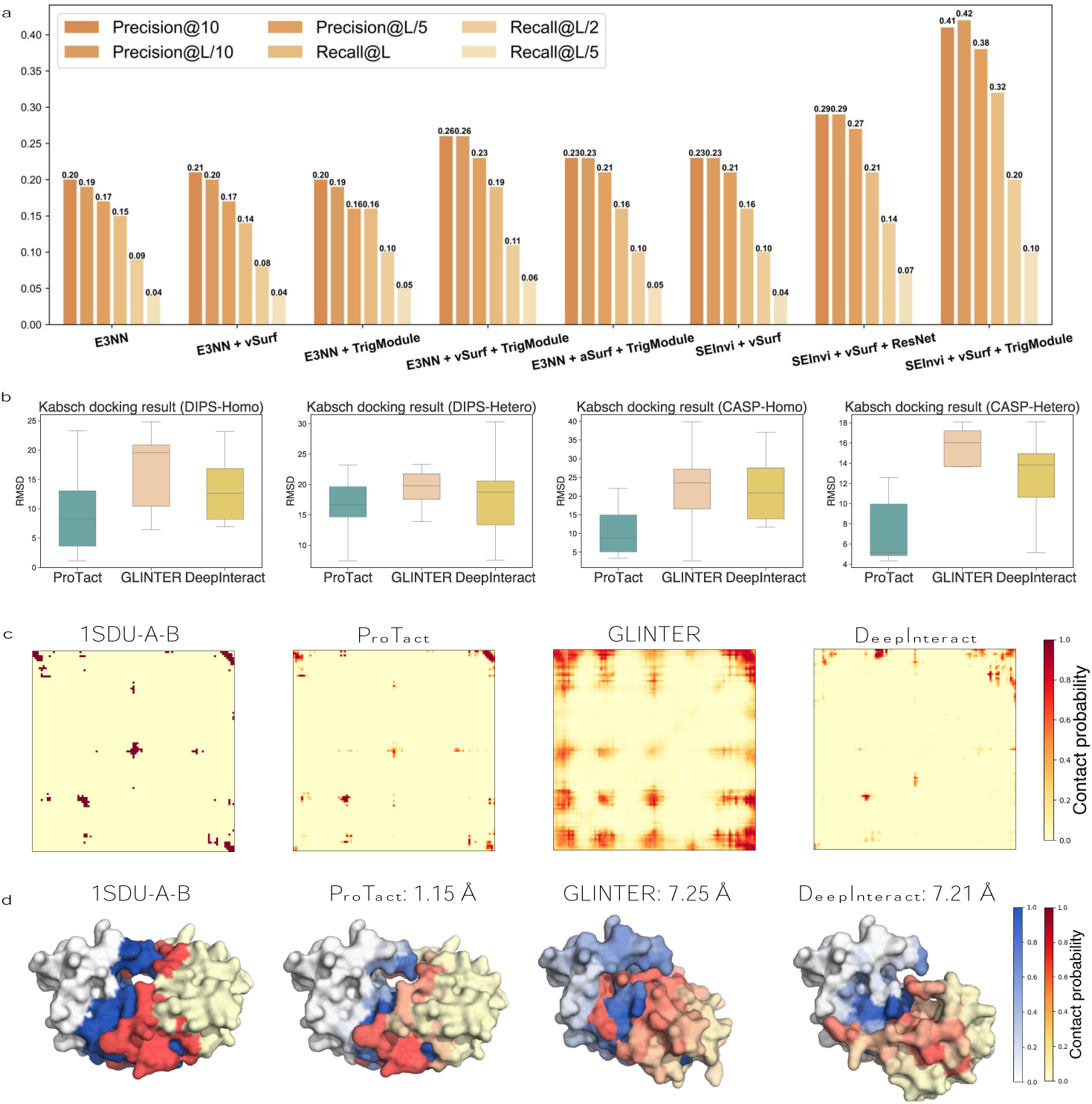
(a) Ablation study of the performance of different configurations of ProTact in predicting protein inter-chain contacts. (b) Performance comparison on the RMSD metric for different protein-protein interaction prediction methods on the DIPS-Plus and CASP13&14 test targets. (c) and (d) Ground-truth and docking predictions for PDB entry 1SDU. For each method, we show the predicted and ground truth interaction map.

We further evaluated ProTact’s superiority by evaluating its impact on docking accuracy with the Kabsch algorithm. The Kabsch algorithm is a well-established method for calculating the optimal rigid-body transformation (rotation and translation) between two sets of points, which is particularly useful in molecular docking to align predicted protein structures with reference structures. By using the Kabsch algorithm, we can observe which predicted contacts remain consistent after alignment, thus validating their effectiveness and reliability. Specifically, ProTact’s docking pose predictions achieved average RMSD values of 9.27 Å (DIPS-Homo), 17.23 Å (DIPS-Hetero), 10.38 Å (CASP-Homo), and 7.39 Å (CASP-Hetero) (Fig. 2b). These results surpassed competing methods such as GLINTER (DIPS: 16.63 Å & 20.20 Å; CASP: 22.68 Å & 15.72 Å) and DeepInteract (DIPS: 13.39 Å & 18.41; CASP: 12.54 Å & 21.89 Å). The poor performance of GLINTER and CDPred could be attributed to their separation of homo- and hetero-interaction predictions. Accurate interface predictions are advantageous for graph-matching algorithms. We further highlighted ProTact’s capabilities by examining two examples (Fig. 2c-d and Fig. S2) from the testing set, underscoring its ability to discern various interfaces, even when overlapping or underrepresented in the PDB. For instance, the HIV protease complex (PDB ID: 1SDU) presented a unique geometry challenging traditional methods, and ProTact accurately predicted both interfaces, including the overlapping regions.

### 2.4 ProTact enhances protein-protein docking performance

We next explored whether ProTact predictions could be used to enhance the modeling of protein complexes. We first aimed to directly improve the performance of docking algorithms by integrating ProTact’s accurate contact predictions as contact restraints. Another validation is provided by demonstrating how ProTact improves the performance of docking algorithms by re-ranking their top-N predictions.

First, we improved HDOCK by using our contact predictions as contact restraints for protein-protein docking. This approach significantly enhanced docking accuracy, as measured by RMSD on the Top-1 predictions, for 74% (37/50) of the tested protein-protein interactions (PPIs) (Fig. S3). As shown in Figure 3a, we also observed an increase in DockQ for 15 out of 17 low docking quality (DockQ *<* 0.23) complexes, including 2G3O-C-E (Fig. 3b), where there was a remarkable improvement from 0.010 to 0.720. These findings emphasize the value of our contact predictions in boosting the performance of established docking methods, particularly for challenging cases like 2G3O-C-E, which had only around 200 multiple sequence alignments (MSAs).

**Fig. 3.**
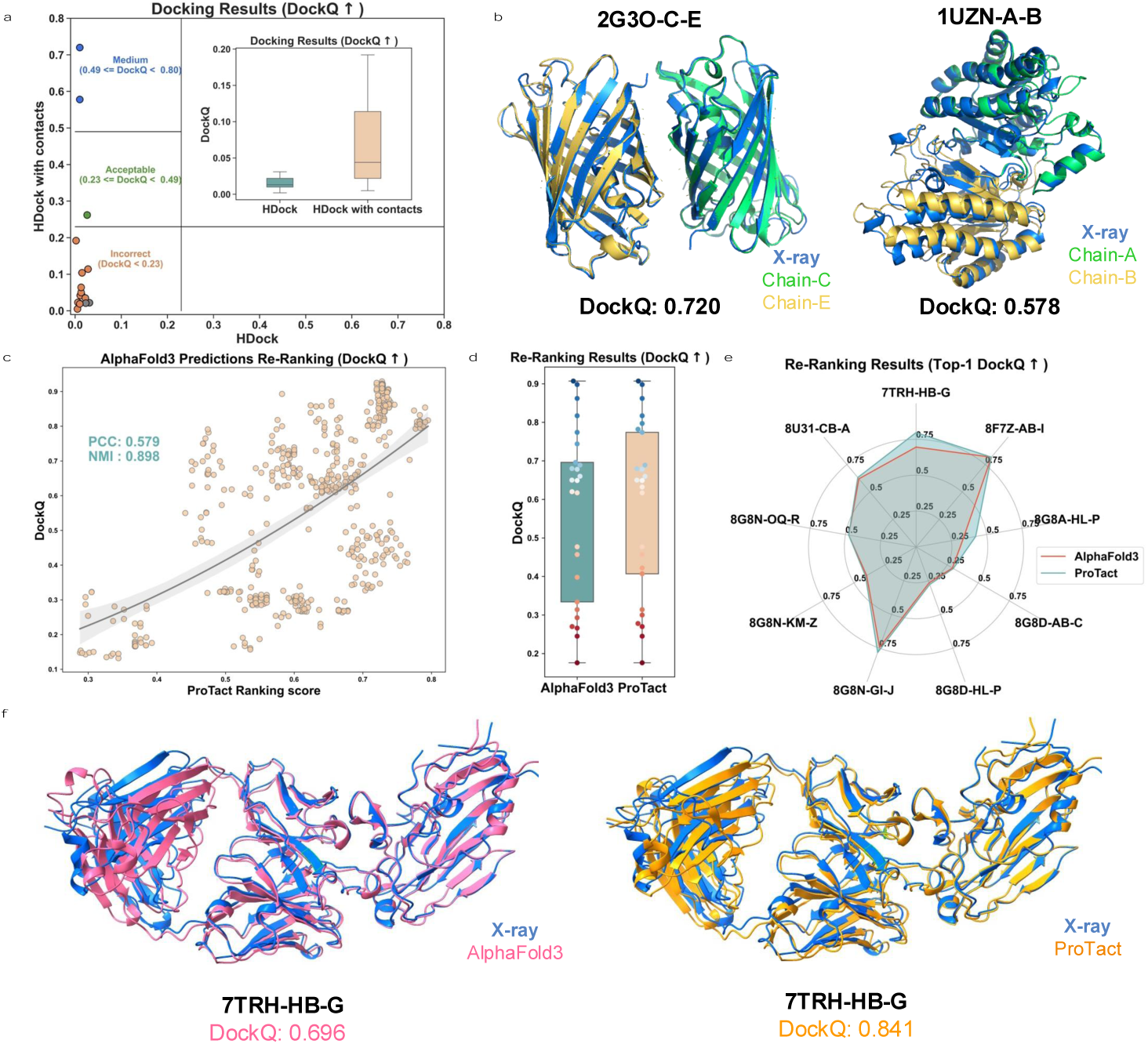
(a) The head-to-head comparison of the top 1 model predicted by HDOCK with and without using ProTact predicted contacts as restraints on the DIPS-Plus and CASP13&14 test targets. (b) are ground-truth and docking predictions with using ProTact predicted contacts for PDB entry 2G3O and 1UZN. (c) ProTact Re-ranking confidence score versus DockQ for antigen-antibody complexes targets. (d) and (e) show the head-to-head comparison of AlphaFold3 and ProTact ranking results. (f) is the representative sample (PDB Entry 7TRH) with AlphaFold3 and ProTact Ranking.

In parallel, we also employed the trained ProTact model as a re-ranking tool for complex structure predictions. We re-ranked the predictions from AlphaFold3 as an example. To achieve this, we focused on antigen-antibody complexes released between January 1st, 2023, and December 31st, 2023, ensuring a maximum sequence identity of 70% and a resolution cutoff of 3.0 Å. The application of the trained ProTact model in conjunction with the ranking score led to notable improvements in the overall docking performance. Specifically, as shown in Fig. 3c, the re-ranked predictions showed a stronger correlation with DockQ scores, achieving a PCC (Pearson Correlation Coefficient) of 0.579 and an NMI (Normalized Mutual Information) of 0.898, compared to AlphaFold3’s PCC of 0.557 and NMI of 0.689 (Fig. S4). This indicates that our ranking score is more closely aligned with the actual docking quality. Furthermore, as shown in Figure 3d-e, among the 25 complexes, the re-ranked Top-1 and Top-5 DockQ values were either equal to or greater than the original AlphaFold3 rankings, with seven complexes experiencing an enhancement in DockQ. Two representative samples, 7TRH-HB-G (Fig. 3f, DockQ from 0.696 to 0.841, Rank 5 to Rank 1) and 8G8A-HL-P (Fig. S5, DockQ from 0.334 to 0.422, Rank 6 to Rank 1) demonstrated significant improvements in DockQ with ProTact re-ranking.

### 2.5 ProTact improves the prediction performance of antigen-antibody complexes

Antibodies are a class of important therapeutic molecules, yet predicting the binding structure of antigen-antibody complexes remains a challenge, primarily due to the lack of MSA information. Our approach, however, shows a notable advantage in such scenarios.

As shown in Figure 4a, we employed a transfer learning strategy to predict inter-chain contacts in antigen-antibody complexes. Figure 4b and Table S4 present the average top-k precision and recall values on antigen-antibody test targets. ProTact consistently outperformed its competitors, especially GLINTER and CDPred, across all metrics. Additionally, our model demonstrated significant improvements in RMSD compared to leading models such as GLINTER and DeepInteract (Figure 4c, Table S5). This superior performance is notable given the scarcity of MSA information (with fewer than 1000 effective sequences, calculated using CD-Hit [36]). This enhancement underscores ProTact’s ability to generate accurate contact map predictions, especially in challenging protein-protein docking scenarios.

**Fig. 4.**
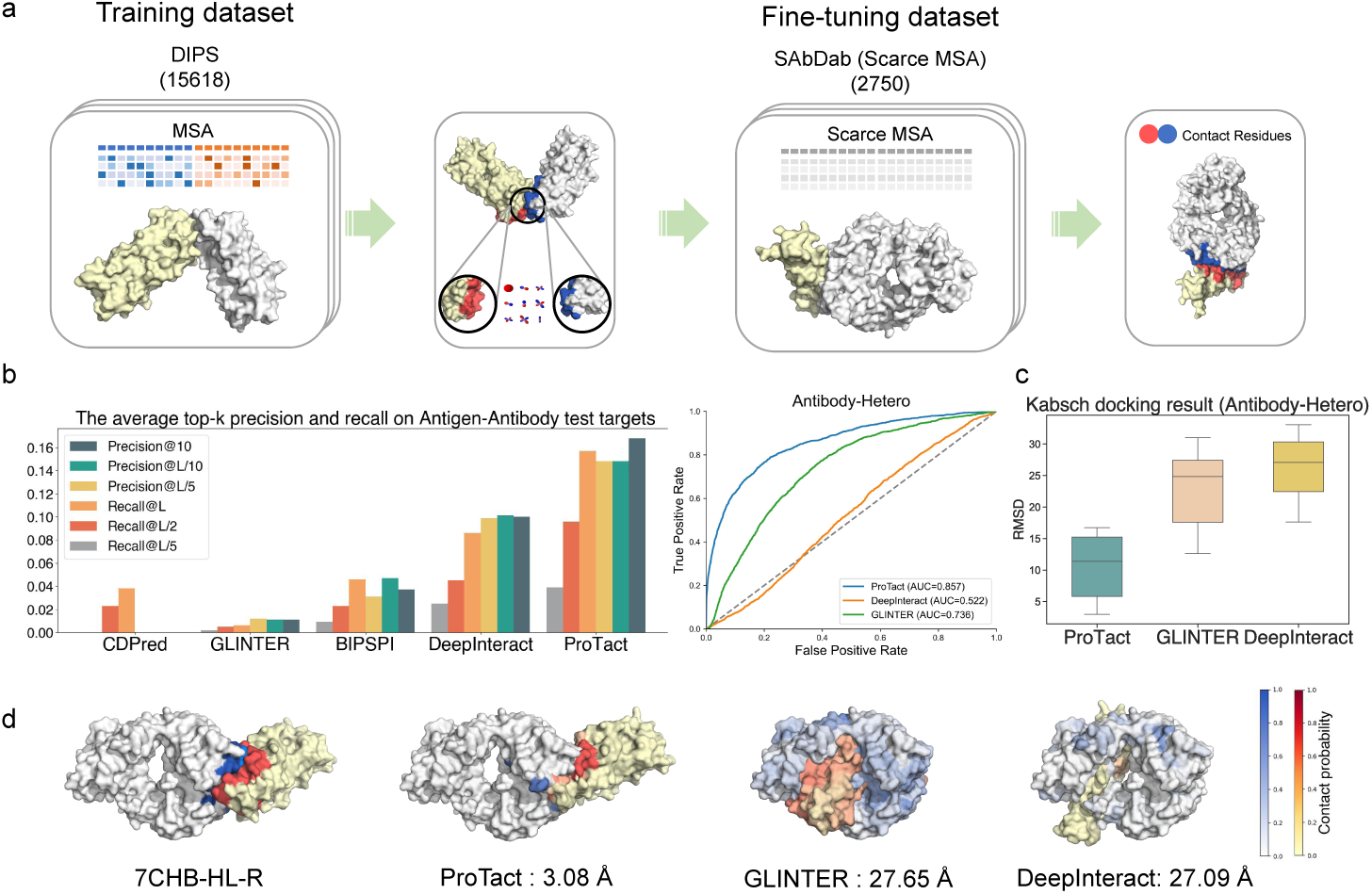
(a) antigen-antibody binding prediction without MSA information using transfer learning and ProTact network. (b) Comparison of the results obtained with the ProTact network to other models on the test set. (c) Performance comparison on binding pose generation combined with the Kabsch algorithm. (d) Case study for 7CHB protein.

As an example, we selected the crystal structure (PDB ID: 7CHB, Figure 4d) of the SARS-CoV-2 Receptor Binding Domain in complex with the BD-236 Fab, providing insights into how this potent neutralizing antibody inhibits SARS-CoV-2. In this case, our model achieved an RMSD of 3.08 Å, significantly outperforming GLINTER (RMSD: 27.65 Å) and DeepInteract (RMSD: 27.09 Å). Analyzing this outcome is important because it represents a challenging case due to the lack of MSA information. Typically, the depth of MSA information affects docking results, with a lack of sequence context often leading to poorer predictions. Despite these challenges, our model delivered the best results among all comparative methods. This success is attributed to our model’s ability to leverage geometric complementarity, providing a more accurate representation of how proteins interact in three-dimensional space. This suggests the robustness and resilience of our ProTact model, even in less-than-ideal circumstances.

## 3 Discussion

In this work, we propose ProTact, a method for predicting inter-chain contacts in protein-protein interactions by integrating trigonometric constraints and molecular surface invariant features. Our extensive evaluations demonstrate that ProTact consistently outperforms state-of-the-art sequence-based and structure-based methods on benchmark datasets, achieving a 31.63% relative improvement in Precision@10 for CASP13&CASP14 targets and a 31.94% relative gain for DIPS-Plus sets. We have also shown that the ProTact framework is not limited to experimental structures. Our experiments demonstrated how ProTact can effectively predict contacts for proteins without available experimental structures, working in conjunction with structure prediction methods.

ProTact significantly enhances the performance of various docking algorithms, including HDock and AlphaFold3. By integrating ProTact’s contact predictions, these algorithms achieve more accurate predictions of protein-protein complexes. Additionally, ProTact’s ability to improve performance is particularly evident in the low-MSA regime, such as antigen-antibody interactions. This enhancement makes ProTact a valuable tool for advancing the accuracy and reliability of protein-protein interaction predictions.

ProTact not only bridges a critical gap in computational biology but also opens up new avenues for improving the accuracy of protein-protein interactions, especially in contexts where evolutionary information is scarce. This advancement paves the way for future research and practical applications in drug discovery, antibody design, and the elucidation of complex biological mechanisms. Despite its strengths, ProTact has limitations that warrant further investigation. One limitation is the incorporation of protein language models and additional geometric and physical constraints. Integrating these elements could further enhance the accuracy and efficiency of ProTact. Additionally, extending ProTact to handle more complex multi-protein systems and dynamic interactions will broaden its applicability and impact in structural biology and drug discovery.

## 4 Methods

### 4.1 Model architecture

Our model consists of two main components: an invariant convolutional encoder (SEInvi) and a trigonometry module (TrigModule). The encoder is responsible for converting protein structure, and sequence features into representations. Then, the interaction module can pass the message of two proteins on an interaction map. Learning the structural feature of the protein is a challenging problem. We aim to obtain a sufficient protein representation from structural and sequence information. This representation must be independent of the coordinate system in which the protein is located; that is, it has rotation and translation invariance. To this end, we propose a SE(3)-invariant encoder named SEInvi based on the E3NN with the vSurf feature to achieve this. For the interaction module, we propose a heterogenous trigonometry network (TrigModule), inspiring from previous works, on the graph structure to capture critical points of the embedding space, as ResNet cannot effectively handle the vector representation generated by SEInvi as well as the intricate many-body effects.

### 4.2 SEInvi: Invariant Convolution module

We upgrade the SE(3) neural network [37] from equivariance to invariance by convoluting spherical harmonics of the dihedral angle and referring to it as SEInvi. The dihedral angle is based on the normal plane of the surface between two residues. First, we input a graph *G* = (*V, E*), where *V* represents the node features and the position of residues and *E* represents the edge features.

#### 4.2.1 Spherical harmonics of dihedral angle

To maintain the invariance of the model, we transform the surface normal vector into the dihedral angle that is calculated based on the direction and two normal vectors of two residues. Formally,

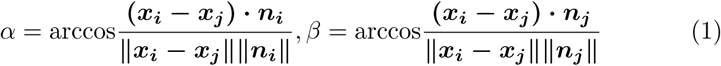

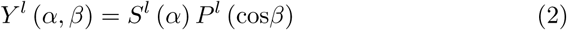

where *α* and *β* are the angles between normal vectors and the direction vector; *S^l^* are the n-dimensional spheres; *P^l^* are the Legendre polynomials. Because the *α* and *β* are invariant when the conformation is rotated or translated, we get the invariant input before the convolution and keep the invariance of the model. Spherical harmonics map the dihedral angle to Hilbert space for convolution, enhancing the model geometry awareness.

#### 4.2.2 Convolution module

The input of Graph *G* containing *V* and *E* will be convoluted on invariant layers. We set the feature configuration in the second spherical harmonics order in the three layers SEInvi. The convolution layer makes the massage pass on the invariant spherical harmonics and aggregate the node and edge features of the neighbours. Formally,

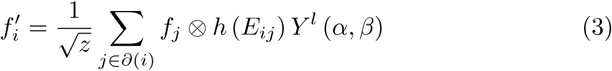

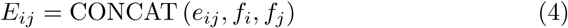

where *f_i_* and *f_j_* denote the node feature of *i* and *j*, *∂* (*i*) denotes the neighbourhoods of the node *i*, *e_ij_*denotes the edge feature between the node i and j, and z represents the degree the node *i* and CONCAT means the concatenation function. We update the edge feature in the last layer of SEInvi by using a multi-layer perceptron on the last *E_ij_*. After convoluting those invariant features, we get the SE(3)-invariant representation containing the node and edge features of the protein.

### 4.3 TrigModule: Trigonometry Module

The trigonometry module updates the interaction map using edge features from both protein graphs. Since the protein graph is incomplete, we initialize the interaction map with node features from both proteins as 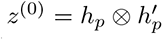 .

Here, *h_p_* and *h*^′^*_p_* represent the node features from the two proteins.

#### 4.3.1 Co-evolutionary features

MSAs are generated for homodimers or heterodimers as input for the calculation of their co-evolutionary features. To challenge the deep learning method to effectively predict inter-chain distance maps from noisy inputs, in the training phase, we use less sensitive tools or smaller sequence databases to generate MSAs, but in the test phase, we use state-of-the-art tools and larger databases to generate the requisite MSAs. Specifically, in the training phase, for a homodimer, we use PSI-BLAST[38] to search the sequence of a monomer against Uniref90 (2018-04) to generate the MSAs, and for a heterodimer, we follow the procedure in FoldDock [39] using the HHblits [40] to search against Uni-clust30 (2017-10) to generate the MSA for each of the two monomers and then pair the two MSAs to produce an MSA for the heterodimer according to the organism taxonomy ID of the sequences.

In the test stage, for a homodimer, we use HHblits to search the sequence of a monomer against the Big Fantastic Database (BFD)[32] and Uniclust30 (2017-10), respectively, to generate two MSAs for a single chain of the homodimers, which are used to generate input features separately to make two predictions that are averaged as the final predicted distance map; for a heterodimer, an MSA is generated by the same procedure used in EvComplex2[33], which applies the jackhammer to search against Uniref90 (2018-04) to generate one MSA for each of the two monomers and then pairs the sequences from the two MSAs to produce an MSA for the heterodimer according to the highest sequence identity with the monomer sequences in each species. The MSA for a homodimer or a heterodimer is used by a statistical optimization tool CCMpred[35] to generate a residue-residue co-evolutionary score matrix (L × L × 1) as features and by a deep learning tool MSA transformer[41] to generate residue-residue relationship (attention) matrices (L × L × 144) as features. L is the number of columns in MSA.

#### 4.3.2 Trigonometry Learning

We implement the trigonometry module for protein-protein interaction prediction by adapting the original method designed for protein-compound interaction. This module is designed to process two protein graphs and update the contact map between them. The trigonometry module ensures that the interaction between two residues from different proteins depends on all edges in both proteins, considering trigonometry and physical constraints such as the excluded volume (Van Der Waals)[42] and saturation effect. Moreover, the trigonometry module can better capture the complementary characteristics between the two interacting proteins, which plays a crucial role in determining the strength and specificity of protein-protein interactions. This improvement in capturing complementary properties leads to more accurate predictions of protein contact maps and, consequently, more reliable protein-protein interaction predictions.

#### 4.3.3 Interaction Learning

The interaction map will be updated by the edge embedding output by SEInvi. First, we apply gated linear transformations for proteins:

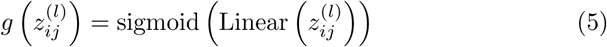

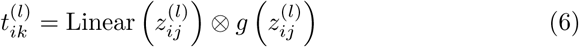

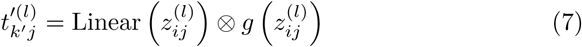

Next, we update the interaction map with these gated linear transformations:

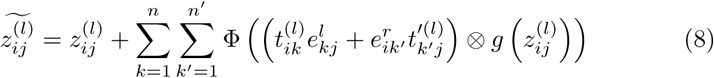

Here, 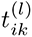 and 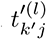 are gated linear transformations of 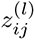 with non-shared parameters, 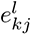 and 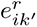 are the edge embedding of the left and right graph, and Φ is a layer normalization function followed by a linear transformation.

#### 4.3.4 Self-attention module

To account for the excluded volume and saturation effects, we design a self-attention module that modulates the interaction between a protein node and all other nodes in the second protein:

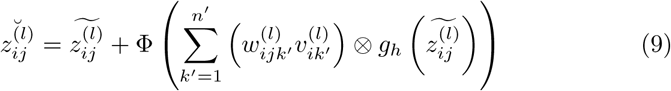

Here, 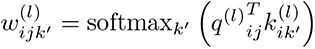 is a set of attention weights, and 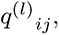 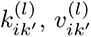 are linear transformations of 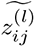

#### 4.3.5 Non-linear transition module

Finally, the interaction embedding is passed through a multilayer perceptron (MLP) to transition to the next layer:

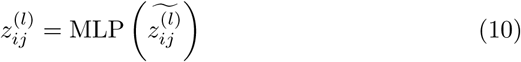

The whole trigonometry module comprises three consecutive parts: the trigonometry update, self-attention modulation, and non-linear transition module. Layer normalization is applied to every input *z*^(*l*)^, and a 25% dropout is applied to the trigonometry update and self-attention modulation during training. The final outputs for protein-protein interaction prediction can be predicted directly based on the last layer embedding *z*^(*L*)^.

### 4.4 Kabsch Algorithm

We employ the Kabsch algorithm [43], a widely used method for the optimal superimposition of two protein structures, to dock the two proteins based on the contact map predicted by ProTact. The Kabsch algorithm computes the optimal rotation matrix that minimizes the root-mean-square deviation (RMSD) between two sets of corresponding points, in this case, the residue coordinates of the two proteins. Here we refer to the algorithm [40] for more details. To improve the docking results and make the surface representation more reasonable, we incorporate MaSIF vectors into the docking process. To do this, we duplicate the contact map for the MaSIF vector based on the position and extend the vector length to 6 Å and Table S4 demonstrates the effectiveness of the MaSIF vector and its contribution to enhancing the docking results obtained by the Kabsch algorithm. This addition helps better to capture the interactions and constraints between the protein surfaces, leading to more accurate protein docking based on the predicted contact map.

### 4.5 AlphaFold3 Re-ranking

We constructed a dataset of antigen-antibody complexes that were generated between January 1st, 2023, and December 31st, 2023. To ensure diversity within our dataset, we imposed a maximum sequence identity threshold of 70% and a resolution cutoff of 3.0 Å to filter the complexes. For each sample in the dataset, we utilized AlphaFold3 to generate 20 predicted structures by varying the seed values. These predictions were then evaluated using a trained ProTact model, which assesses the quality of the predicted contact maps.

The ProTact model takes the predicted structures as input and outputs inter-chain contacts with additional information on multi-body interactions and surface information. We compared the ProTact predictions against the original AlphaFold3 predictions to evaluate their accuracy and precision. The evaluation scores from ProTact were combined with the original ranking scores provided by AlphaFold3 for each predicted structure. This integration of scores allowed us to produce a new ranking of the predictions, prioritizing those with higher confidence in both structural accuracy. Through this method, we aimed to refine the prediction rankings and select the most reliable models for further analysis or application.

### 4.6 Datasets

We trained ProTact on DIPS-Plus, the largest feature-rich dataset of protein complexes for protein interface contact prediction. We tested our model on the test dataset of DIPS-Plus and CASP13&14, respectively. In the scarce MSA scenario, we fine-tuned and tested our model on the antibody-antigen benchmark SAbDab.

#### DIPS-Plus

DIPS is a large protein complex structures dataset mined from the Protein Data Bank and tailored for rigid body docking. Following [24] we select 32 homodimers and heterodimers to compare the performance of our model. After removing proteins with ¿= 30% sequence identity with the test datasets, 15,618 and 3,548 binary complexes are left for training and validation. 80% of these complexes are randomly selected for training while 20% are for validation.

#### CASP13&14

We also evaluate each method on 14 homodimers and 5 heterodimers with PDB structures publicly available from the 13th and 14th sessions of CASP-CAPRI as these targets are considered by the bioinformatics community to be challenging for existing interface predictors as Morehead et al. For any CASP-CAPRI test complexes derived from multimers (i.e., protein complexes that can contain more than two chains), we chose the pair of chains with the largest number of interface contacts to represent the complex.

#### SAbDab

SAbDab is a database containing all the antibody structures available in the PDB. We split the antigen-antibody structure into two parts: the antibody containing the heavy and light chains and the antigen. We follow the setting of Luo et al.(2022)24, leaving 19 antibody-antigen complexes for testing and 2705 antibody-antigen complexes which are L1 L2*<*65536 for training (L1 and L2 represent the length of antibody and antigen).

#### BlindTest

To create a larger blind test dataset, we collected the heterodimers and homodimers published between 2023-01-01 and 2023-06-31 in the PDB. After filtering out similar sequences at a sequence identity threshold of 40% and excluding sequences with *>*1000 residue targets, 191 heterodimers and 41 homodimers were selected to create a larger blind test set. Herein, we utilized a training set similar to that of DeepInter, consisting of 3,504 homomeric PPIs and 2,174 heteromeric PPIs, with a sequence identity threshold of 40% and a distance cutoff of 8 Å.

### 4.7 Evaluation

To evaluate the performance of ProTact, we follow the setting established by [24], where a positive label is assigned to each inter-chain residue pair found within 6/8 Å of each other in the complex’s bound (i.e., structurally-conformed) state. We use several evaluation metrics to assess the accuracy and robustness of our method in predicting protein-protein interactions and the quality of the generated docking results.

#### 4.7.1 Average top-k precision (P@k)

P@k measures the percentage of correct inter-chain residue pair predictions among the top k predictions. It evaluates the model’s ability to predict relevant contacts accurately.

#### 4.7.2 Top-k Recall (R@k)

R@k measures the proportion of true positive interactions among the top k predictions. It evaluates the model’s ability to identify true positive inter-chain residue pairs.

#### 4.7.3 Receiver Operating Characteristic Area Under the Curve (ROC-AUC)

ROC-AUC is used to measure the model’s ability to discriminate between positive and negative inter-chain residue pairs. A higher AUC indicates better performance in distinguishing true positive interactions from false positives.

#### 4.7.4 Root Mean Square Deviation (RMSD)

RMSD measures the average distance between the atoms of the superimposed structures, reflecting the accuracy of the predicted protein-protein docking results. In this article, we use the unit of RMSD as Å. Lower RMSD values indicate better alignment between the predicted and experimental structures, signifying more accurate docking predictions. This study applies the Kabsch algorithm to the predicted contact maps to perform protein-protein docking and calculate the RMSD between the predicted and experimental structures.

#### 4.7.5 DockQ

DockQ [44], a continuous protein-protein docking model quality measure derived by combining *F_nat_*, LRMS, and iRMS to a single score in the range [0, 1] that can be used to assess the quality of protein docking models.

### 4.8 Implementation and training

ProTact is trained on one A800 graphics processing unit (GPU) with PyTorch (v1.8.0) and the optimizer AdamW. In addition, the training process uses a learning rate of 0.001 without any regularization and a dropout of 0.1 for the triangle-aware module. Because of the large use of GPU memory by the triangle self-attention layer, we apply a mini-batch size of 1 and set a maximum length of 256 for each monomer sequence in the training process. Specifically, for the monomer proteins of the dimer with a sequence length of more than 256, we use a window of size 256 and a stride of 1 to scan the sequence. Fragments with the maximum inter-protein contacts are kept, and the final cropped sequences are randomly selected from those fragments to represent the protein sequence. During training, the ground-truth inter-protein contacts are defined as those inter-protein residue–residue pairs with a distance of *<* 6 Å. Here, the distance between two residues is represented by the minimal distance between the heavy atoms of the two residues.

## Data availability

We used freely available data as described in Methods. The data and code to reproduce the datasets and experiments are available at https://github.com/biomed-AI/ProTact.

## Acknowledgements

This study has been supported by the National Key R&D Program of China [2020YFB0204803], National Natural Science Foundation of China [62041209, 62402314, 12126610], Guangdong Key Field R&D Plan [2019B020228001, 2018B010109006], and Guangzhou S&T Research Plan[202007030010, 202002020047]. S. Z. acknowledges funding from the Asian Young Scientist Fellowship.

## Competing Interests

The authors declare that no competing interests exist.

## Authors’ contributions

S.Z. and Y.Y conceived and supervised the project. J.R., D.L., S.Z., and J.Z. contributed to the algorithm implementation. J.R.,D.L. and S.Z. contributed to the visualization and server implementation.J.R., D.L., S.Z. and Y.Y wrote the manuscript. All authors were involved in the discussion and proofread.

## Notes

### Competing Interest Statement

The authors have declared no competing interest.

